# Cell Maps for Artificial Intelligence: AI-Ready Maps of Human Cell Architecture from Disease-Relevant Cell Lines

**DOI:** 10.1101/2024.05.21.589311

**Authors:** Timothy Clark, Jillian Mohan, Leah Schaffer, Kirsten Obernier, Sadnan Al Manir, Christopher P Churas, Amir Dailamy, Yesh Doctor, Antoine Forget, Jan Niklas Hansen, Mengzhou Hu, Joanna Lenkiewicz, Maxwell Adam Levinson, Charlotte Marquez, Sami Nourreddine, Justin Niestroy, Dexter Pratt, Gege Qian, Swathi Thaker, Jean-Christophe Bélisle-Pipon, Cynthia Brandt, Jake Chen, Ying Ding, Samah Fodeh, Nevan Krogan, Emma Lundberg, Prashant Mali, Pamela Payne-Foster, Sarah Ratcliffe, Vardit Ravitsky, Andrej Sali, Wade Schulz, Trey Ideker

## Abstract

This article describes the Cell Maps for Artificial Intelligence (CM4AI) project and its goals, methods, standards, current datasets, software tools, status, and future directions. CM4AI is the *Functional Genomics Data Generation Project* in the U.S. National Institute of Health’s (NIH) *Bridge2AI* program. Its overarching mission is to produce ethical, AI-ready datasets of cell architecture, inferred from multimodal data collected for human cell lines, to enable transformative biomedical AI research.

## Introduction

### 1. Goals of CM4AI

Cell Maps for Artificial Intelligence (CM4AI) is an NIH-funded Bridge to Artificial Intelligence (Bridge2AI)^1^ Data Generation Project (DGP) in the area of Functional Genomics. It consists of three pillars – Data, People, and Ethics – organized into six modules (Data: Data Acquisition, Tools, and Standards; People: Skills and Workforce Development, and Teaming; and Ethics).

CM4AI’s objective is to deliver machine-readable hierarchical maps of cell architecture as AI-Ready data produced from multimodal interrogation of 100 chromatin modifiers and 100 metabolic enzymes involved in cancer, neuropsychiatric, and cardiac disorders in disease-relevant cell lines under perturbed and unperturbed conditions, utilizing state-of-the-art mass spectrometry based proteomics, spatial proteomics / cell imaging, and genetic perturbations using CRISPR. The cell lines currently under investigation in CM4AI are treated (with paclitaxel and vorinostat) versus untreated MDA-MB-468 breast cancer cell lines; and undifferentiated vs. neurons and cardiomyocytes generated from NIH-iPCS-1 KOLF2.1J induced pluripotent stem cells (IPSCs). CM4AI’s software pipeline and AI-readiness framework provide important and reusable enabling capabilities for this work.

CM4AI input data streams are generated using immunofluorescence (IF) subcellular microscopy for spatial proteomics data; affinity purification mass spectroscopy (AP-MS) and size exclusion mass spectroscopy (SEC-MS) methods for protein-protein interaction (PPI) data; and single-cell CRISPR-Cas perturbation screens by cell type. Input data streams are integrated via the multi-scale integrated cell (MuSIC) software pipeline employing deep learning models and community detection algorithms^2^, and output cell maps are packaged with provenance graphs and rich metadata as AI-Ready datasets in RO-Crate format^3,4^ using an extended, client-server version of the FAIRSCAPE framework^5^.

At a strategic level, CM4AI – as do other Bridge2AI DGPs – aims to ethically leverage the most state-of-the art computational and experimental tools in biomedical science to enable a new generation of transformative artificial intelligence research to benefit humanity. Its datasets and software are available to the research community under minimally restrictive terms consistent with ethical guidelines, researcher attribution, and research integrity.

### 2. What is a Cell Map?

Cell maps are hierarchical directed acyclic graphs (DAG), where each node represents an assembly of proteins in proximity at a given scale, spanning from large assemblies representing cell compartments (e.g. nucleus, mitochondria) to small assemblies of proteins in close proximity (e.g. protein complexes or subunits). The data streams for constructing these maps are generated from a panel of perturbed and unperturbed cell lines, including treated and untreated cancer cell lines as well as differentiated and naive induced pluripotent stem cells (iPSCs). Cell maps provide a foundation for downstream applications in human genomics, including interpretation of genetic variants and mutations. As an example, they have been used in AI tools for “visible machine learning” or “visible neural networks” (VNNs) to interrogate how protein assemblies in the cell affect cell-level phenotype predictions^6–10^.

In the CM4AI cell mapping process, affinity purification-mass spectrometry (AP-MS) and size-exclusion mass spectroscopy (SEC-MS) techniques generate protein interaction networks, while immunofluorescence (IF) staining and high-resolution microscopy reveal protein localization and distribution within human cells^11^.

Cell maps are produced by integrating these localization and interaction data using self-supervised embedding approaches from deep learning, followed by algorithmic community detection at multiple resolutions to generate a hierarchy of protein assemblies^12^. Cell maps are shared via the Network Data Exchange (NDEx)^13^ and can be visualized in a web browser or accessed via tools such as Cytoscape^14^, HiView^15^, and the Python ndex2 library^16^.

An example cell map from the Multi-Scale Integrated Cell (MuSIC)^2^ data set visualized in HiView is shown in Figure 1.

**Figure 1.**
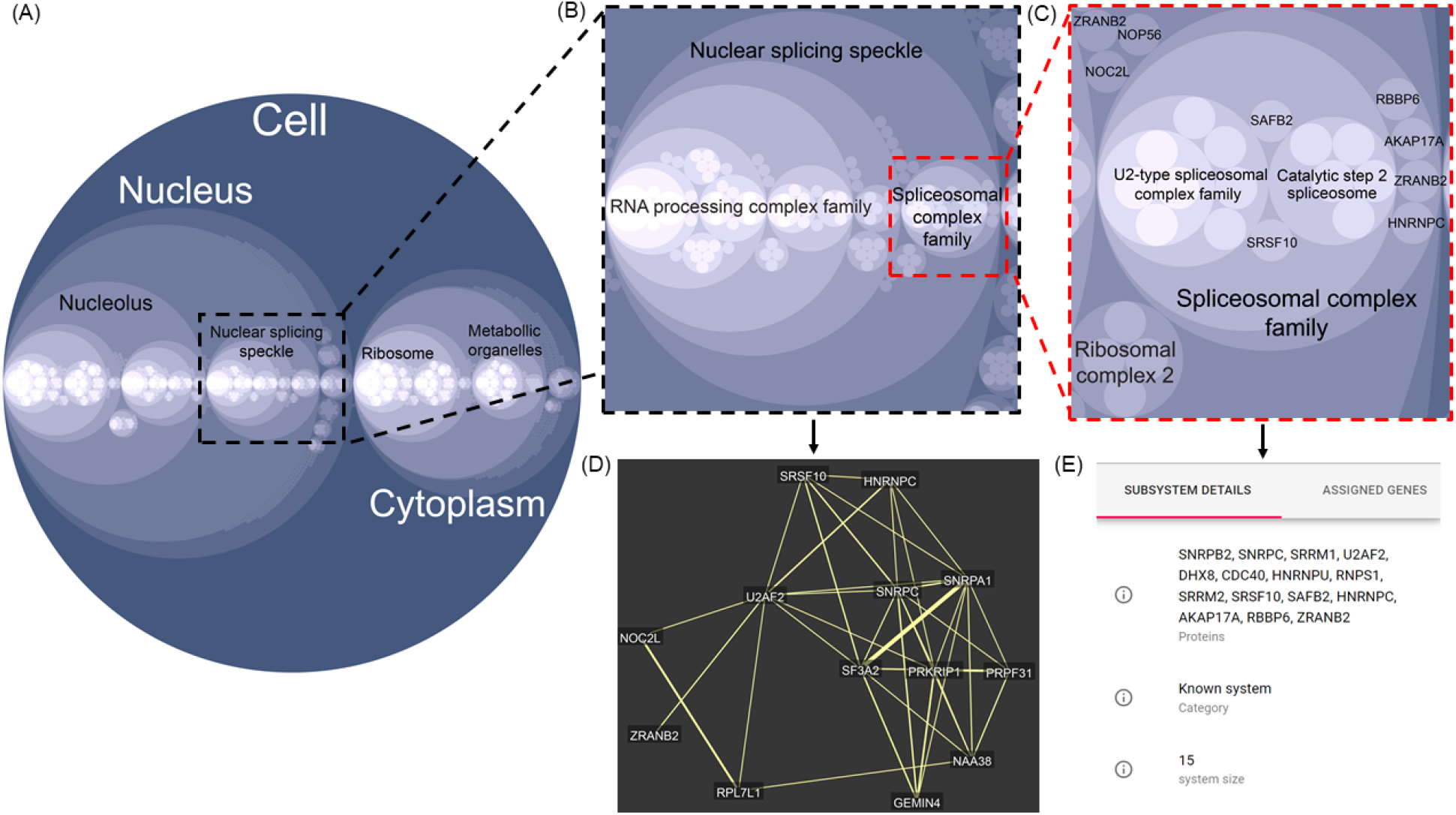
HiView visualization of a hierarchical cell map as a circle-packing diagram. (A) A unified cell map of the interaction network in HEK293 cells generated by integrating immunofluorescence images and affinity purification interactions for 661 proteins.^2^ The hierarchical cell map contains 10 layers of depth representing communities at multiple resolutions, such that communities at smaller distance (lighter circles) are nested inside larger communities (darker circles) similar to physical compartments within a cell (the “root” structure). HiView can be used to zoom in on nested communities, such as (B) the nuclear splicing speckle (depth layer 4) and (C) the spliceosomal complex family (depth layer 5), to begin to resolve smaller interaction communities and even individual proteins. For each community, HiView provides the underlying interaction data that support it. For example, (D) shows the protein-protein interaction network within the nuclear splicing speckle community, where yellow edges represent similarities in the embedding from both protein image and affinity purification data. HiView also provides a list of proteins within each community. For example, (E) shows the list of proteins (15 total) located within the spliceosomal complex family.

### 3. What are Ethical, FAIR, AI-Ready Biomedical Data?

As defined by CM4AI, AI-Ready biomedical data are fully characterized FAIR data of known provenance, which can be ethically and reliably processed by AI applications; whose models and software are available and well-described for validation and re-use; and whose predictions may be fully explained and interpreted to the user as needed. CM4AI data are distinctive within Bridge2AI in that they are non-clinical (from tissue cultures) and are considered to be de-identified as they cannot be matched, with current knowledge, to a human subject.

AI-Readiness has been defined in multiple ways in the literature. Regardless of the specific definitions adopted, in all cases significant requirements are placed on the datasets used to train AI/ML models and on datasets analyzed using these AI/ML models. Datasets that meet these requirements are called “AI-Ready”. We define the following criteria for purposes of this discussion:

⍰ **FAIR** – Datasets, and the software used to prepare them for AI/ML analysis, must comply with the FAIR Principles which have been outlined for data and for software; interoperability is a particular requirement of AI-Readiness^17^. FAIRness is an NIH requirement and is consistent with best current scholarly practice.
⍰ **Provenanced –** Provenance Graphs of computations, data, software, and models used to prepare the dataset must be available as metadata^18–20^.
⍰ **Characterized** – Complete schemas, validation procedures, and data sheets^21^ for all datasets in the provenance graph; and model cards^22^ for all models used to prepare the dataset, are resolvable from the dataset’s metadata.
⍰ **Explainable** – Dataset must be fully provenanced (see above) and statistically characterized within any limitations and constraints, including ethical considerations and limitations; and this characterization is available in the metadata^23–26^;
⍰ **Ethical** – The dataset was obtained, validated, documented, licensed, and distributed ethically^27^. Ethical conditions will include but are not limited to: ethical treatment of subjects including protection of human subjects data and ethical treatment of animals; proper non-biasing data analysis and scientific integrity of methods, conclusions, and publications; full documentation of scientific methods and reagents; and licensing and distribution of data, software, and models with openness to the scientific community for reuse consistent with protection of human subjects privacy and adhering to responsible research conduct; moreover, this requirement involves anticipating potential applications of the data and derived AI/ML systems, integrating them into data governance frameworks to optimize benefits and mitigate potential risks, and promoting their use for the collective good.

These criteria are interdependent. For example, ethical validation capability depends upon explainability, explainability depends upon provenance, and so forth.

CM4AI uses an expanded version of the FAIRSCAPE framework to establish a basis for AI-readiness. As we progress through the project, our AI-readiness features will become ever more complete. At present they comprise: (a) establishing FAIRness including rich metadata and persistent, globally unique identifiers; (b) computing a machine-readable provenance graph, resolvable as metadata for all results, including inputs, computations, software, and outputs; and (c) characterizing and validating all datasets and data elements, and mapping data elements to public ontology vocabularies, where appropriate, using JSON-Schema mini-data-dictionary descriptions resolvable from the provenance graph metadata. Further AI-Readiness capabilities on our near-term research agenda are described in the Methods section.

## Methods

### 1. Cell Lines

CM4AI has elected to use the cancer cell line MDA-MB-468^28^ (+/-treatment with paclitaxel and vorinostat) and the iPSC line KOLF2.1J^29^ (+/-neuronal and cardiomyocytic differentiation), both of which have been ethically sourced.

MDA-MB-468 (RRID:CVCL_0419) is a triple negative breast cancer cell line established from a metastatic site pleural effusion of a 51-year-old black female with a metastatic mammary adenocarcinoma, available from ATCC. This cell line has been extensively used to study triple-negative breast cancer and is well characterized with data such as transcriptomic, mutational profile and whole-genome sequencing available.

The KOLF2.1J (RRID:CVCL_B5P3) iPSCs cell line was derived from a healthy male Northern European donor, available from the Human Induced Pluripotent Stem Cells Initiative (HipSci) resource^30^. It is available for access by non-for-profit organizations via a simple MTA.

### 2. Data Acquisition

#### 2.1 Protein-Protein Interaction

To map protein-protein interactions (PPIs) of 100 chromatin regulators under different conditions (cancer: no treatment, paclitaxel or vorinostat; iPSC: undifferentiated, neurons and cardiomyocytes), we employed AP-MS on endogenously tagged cell lines and size-exclusion chromatography coupled to mass spectrometry (SEC-MS) for proteome-wide complex/interaction mapping as two orthogonal mass spectrometry-based approaches. We have endogenously tagged 17 genes in MDA-MB-468 and acquired AP-MS data under three conditions (untreated, paclitaxel and vorinostat treated) . We are currently in the process of tagging 34 additional genes inMDA-MB-468 cells . In addition, we have performed SEC-MS on MDA-MB-468 cells under three conditions (untreated, paclitaxel and vorinostat treated), which enabled us to identify 72/100 chromatin modifiers, with 52 of them being integral components of protein complexes. In addition to chromatin modifiers of interest, SEC-MS allowed us to map complexes in MDA-MB-468 cells and investigate the impact of treatment proteome-wide. We detected PPI profiles of over a thousand complexes in MDA-MB468 cells, with thousands of proteins exhibiting differential elution profiles between control and paclitaxel or vorinostat treated cells. SEC-MS has also been performed in KOLF2 iPSCs and derivatives, uncovering more than 700 protein complexes in both parental iPSCs and differentiated neurons.

#### 2.2 Spatial proteomics mapping

For Year 1, we proposed to map the spatial subcellular organization of key chromatin modifiers, their interactors and key signaling molecules involved in cancer using the Human Protein Atlas resource of antibodies. We established automated fixation and permeabilization protocols for the pipetting robot for MDA-MB-468 and KOLF2 cell lines, and completed spatial proteomics mapping of 100 chromatin regulators in the MDA-MB-468 cells (+/-paclitaxel or vorinostat), with another 500 proteins pending significant hits from the genetic perturbations and PPI results. The first set of images was released as input for the CM4AI Tools Module’s MuSIC pipeline and processed to RO-Crate outputs.

#### 2.3 Genetic perturbation mapping

For Year 1, we performed single-cell CRISPR screens perturbing 100 chromatin regulators in the MDA-MB-468 cells under 3 conditions (no treatment, paclitaxel or vorinostat) and KOLF2 iPSC in the undifferentiated state. We have designed a CRISPR lentiviral library targeting 100 chromatin factors with 6 guide RNAs per gene. We generated and characterized MDA-MB-468 and KOLF2 CRISPR lines expressing inducible dCas9. Single-cell CRISPR screens of 100 chromatin regulators (+/– drug) in MDA-MB-468 cells and in undifferentiated KOLF2.1J iPSCs were done using the 10x Genomics 3’HT kit. The resulting data are currently being QCed.

### 3. Tools: Cell Map Data Integration Pipeline and VNN Tools

The Tools Module of CM4AI is responsible for producing and maintaining a dataset integration and map production system, the Multi-Scale Integrated Cell (MuSIC) pipeline, which produces integrated cell maps from the multiple, multi-modal input data streams. The MuSIC pipeline is divided into multiple segments, each of which calls the FAIRSCAPE client package to validate inputs and create the output RO-Crate package for that segment which, in turn, is ready for dispatch to the FAIRSCAPE server for PID creation and provenance graph entailment computation. FAIRSCAPE-generated PIDs resolve to complete human- and machine-readable metadata including licensing information, and provide links to the underlying datasets.

MuSIC pipeline segments and their functions are shown below and in Figure 2. The steps are:

**Figure 2.**
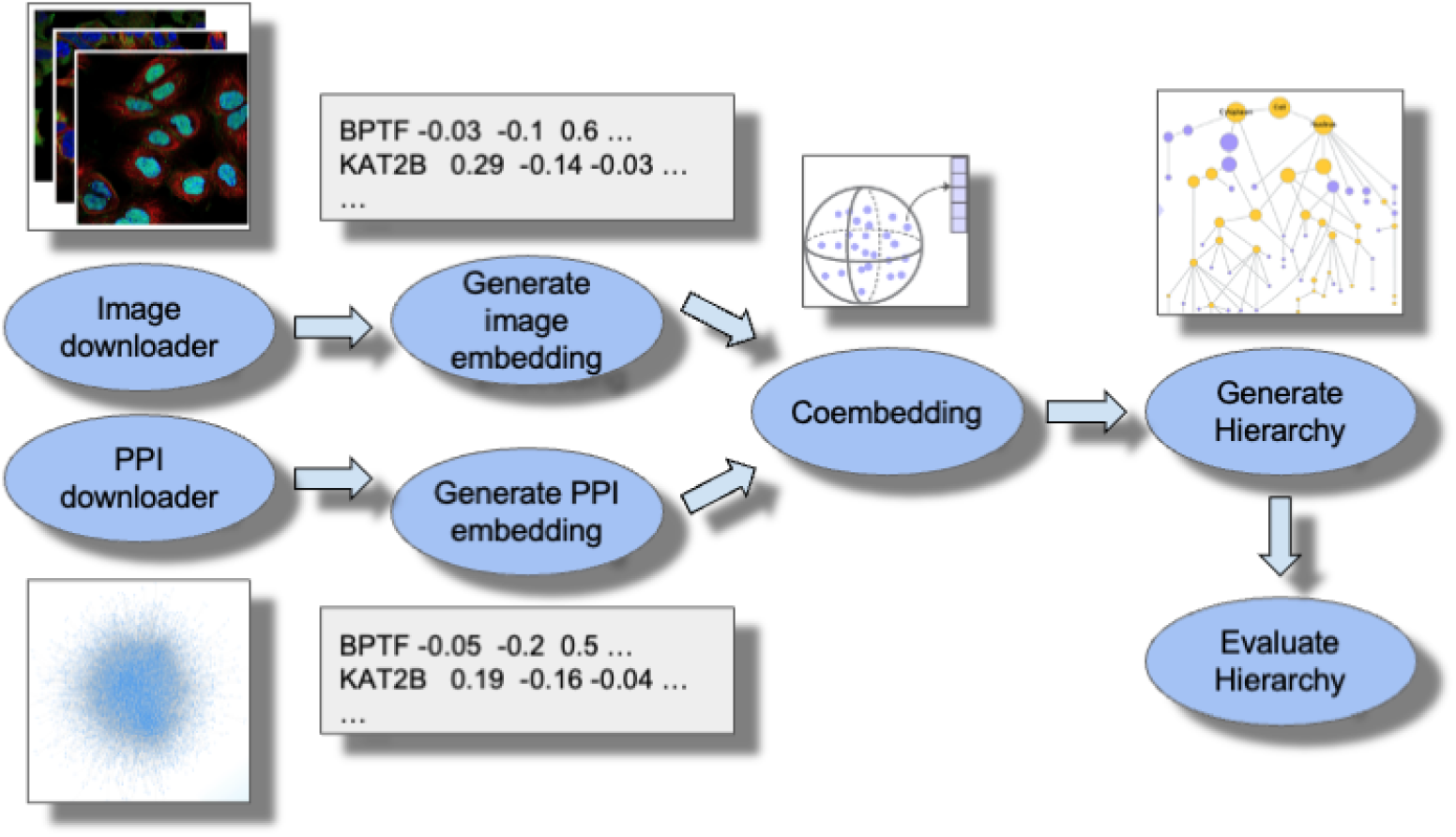
Tools Module: Data Integration and Map Production Pipeline. With the Standards and Data Acquisition Modules, we developed an alpha release of software that interfaces the MuSIC pipeline with the FAIRSCAPE infrastructure and data repository. The system includes data dictionaries, formatting standards, and a FAIRSCAPE metadata and provenance API, implemented as a command line software package.

a) Download PPI and Image Data

-PPI networks for the cell line and conditions are downloaded from the Krogan Laboratory’s deposition archive and made available for further processing.

-IF images for the cell line and conditions are downloaded from the Lundberg Laboratory’s deposition archive and made available for further processing.

b) Generate embeddings

-Downloaded PPI networks are processed using the node2vec deep learning model^31^ to reduce dimensionality, producing a PPI embedding that contains information about the protein’s interactions.

-IF images are processed using a Human Protein Atlas deep learning model^32^ to reduce dimensionality, producing an image embedding that contains information about protein localization.

c) Co-Embedding

-The PPI and image embeddings are integrated to obtain a co-embedding for each protein using contrastive deep learning. This integration model learns co-embeddings such that the original embeddings can be reconstructed with minimal information loss and proteins with similar PPI and image embeddings have similar co-embeddings^33^.

d) Protein Community Detection and Hierarchy Creation

-Community detection is performed based on the all-by-all similarities of pairs of proteins in the co-embedding space. The resulting hierarchy of protein communities is output as the final cell map.

e) Hierarchy evaluation

-Cell maps are annotated in a two pronged approach. First, cell maps are aligned to known protein function and pathway resources, including the Gene Ontology (GO)^34^ and Reactome^35^, to determine protein assemblies in the cell map with high overlap with known cell biology. Second, assemblies in the cell maps are annotated using a large language model (LLM) approach that we developed to name sets of proteins and assign a name confidence score.

f) Integrative structure modeling

We are currently exploring the feasibility of integrative structure modeling of the MuSIC communities. The resulting structural models will increase our understanding of the MuSIC communities and help with planning future experiments.

To begin, we developed a bioinformatics pipeline for annotating MuSIC communities by the available structural information about the community members and their interactions. This structural information includes data from the Protein Data Bank (PDB)^36^, AlphaFold Protein Structure Database (AlphaFoldDB)^37^, crosslinking mass spectrometry, and prediction of disordered sequence segments. We then ranked the MuSIC communities by the amount of available structural information, serving as a proxy for the feasibility of integrative structure modeling. Finally, we performed integrative structure modeling of a few top ranked communities. This process uses our standard integrative structure modeling framework, which proceeds through the following four stages^38–40^: (i) gathering input information, (ii) representing subunits and translating data into spatial restraints, (iii) configurational sampling to produce an ensemble of models that satisfy the restraints, and (iv) analyzing and validating input information and models. The modeling protocol was scripted using the Python Modeling Interface package, a library for modeling macromolecular complexes based on our open-source Integrative Modeling Platform (IMP) package version 2.18 (https://integrativemodeling.org).

### 4. Standards: AI-Readiness Packaging

The Standards Module of CM4AI is responsible for AI-Readiness packaging of data, software, metadata and provenance graphs. This task is performed by the FAIRSCAPE framework. The integration of FAIRSCAPE with CM4AI Data Acquisition and the Tools data integration pipeline is shown in Figure 3.

**Figure 3.**
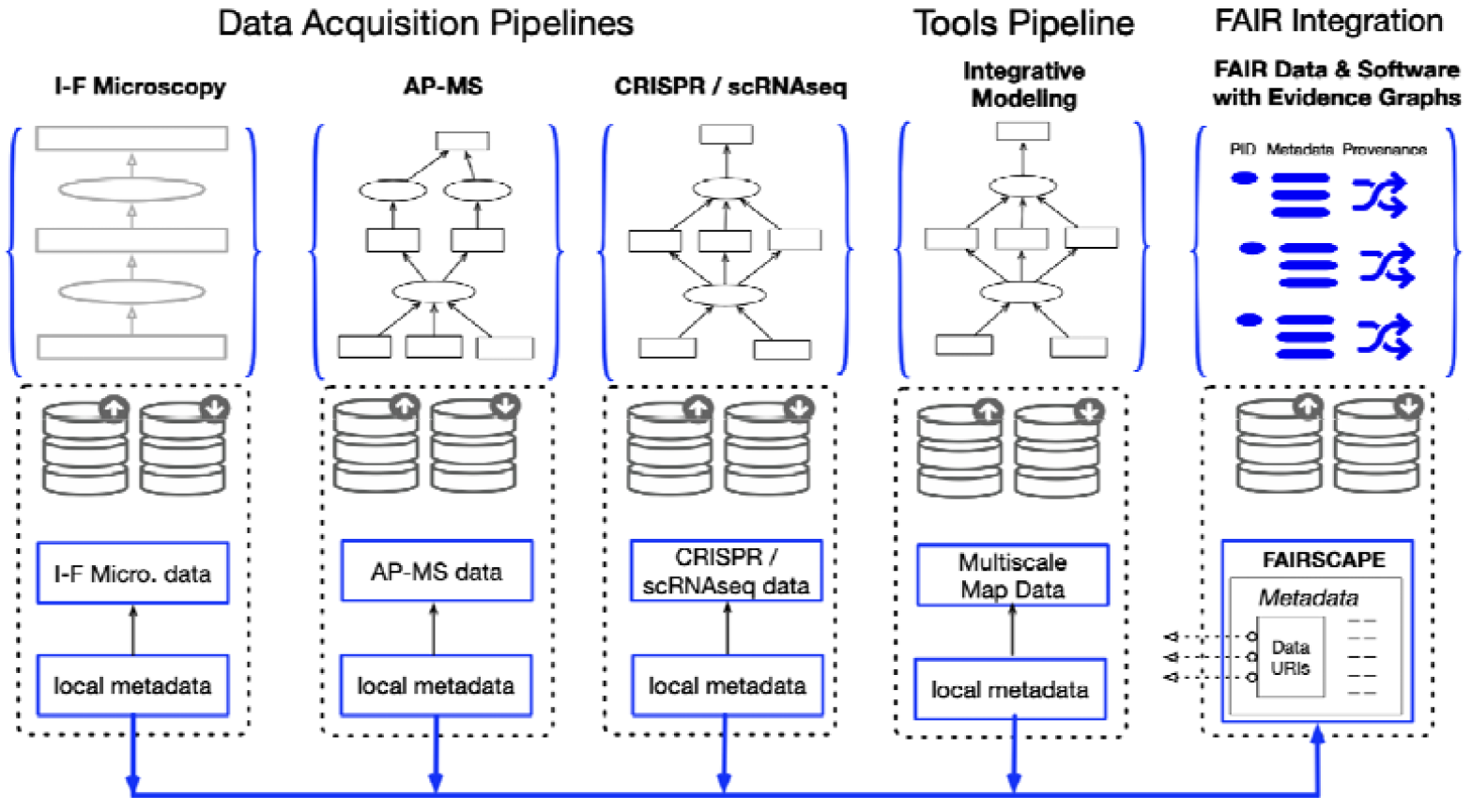
FAIRSCAPE AI-Readiness framework as applied to CM4AI datasets. FAIRSCAPE consists of a client-side Python3 application, called either from the command line or as a set of Python functions by the Tools Module’s data integration pipeline, and a server application, also in Python3, which completes the packaging.

The client-side package, FAIRSCAPE-CLI, is called when any computation or coherent set of computations in the pipeline is completed, and it is passed metadata which defines schemas in JSON-Schema^41^ for the datasets in the computational unit, as well as the inputs, computations, software, models, and outputs. FAIRSCAPE-CLI creates an RO-Crate package with the datasets, metadata, and software – or resolvable references to these components – and unique stubs for identifier creation on each of these components.

The RO-Crates are then sent to the FAIRSCAPE server where they are registered and assigned persistent, resolvable, globally unique persistent identifiers (PIDs). The RO-Crates are then decomposed into their individual components – datasets, models, software – which are also registered and assigned PIDs. The PID system currently in use is the ARK scheme^42^ – with DOIs a future feature as supplementary PIDs for final-state publishable work.

Lastly, the server computes end-to-end entailments on each RO-Crate’s provenance as expressed in the EVI Evidence Graph Ontology and links them together where possible. A graphical view of an evidence graph presented as part of the human-readable landing page for a CM4AI RO-Crate package, is shown in Figure 4. Alternate views serialized in JSON-LD and in RDF-XML are also provided on the landing page. The complete RO-Crate is provided as Supplemental Data, and all RO-Crates for the current release are available in the University of

**Figure 4.**
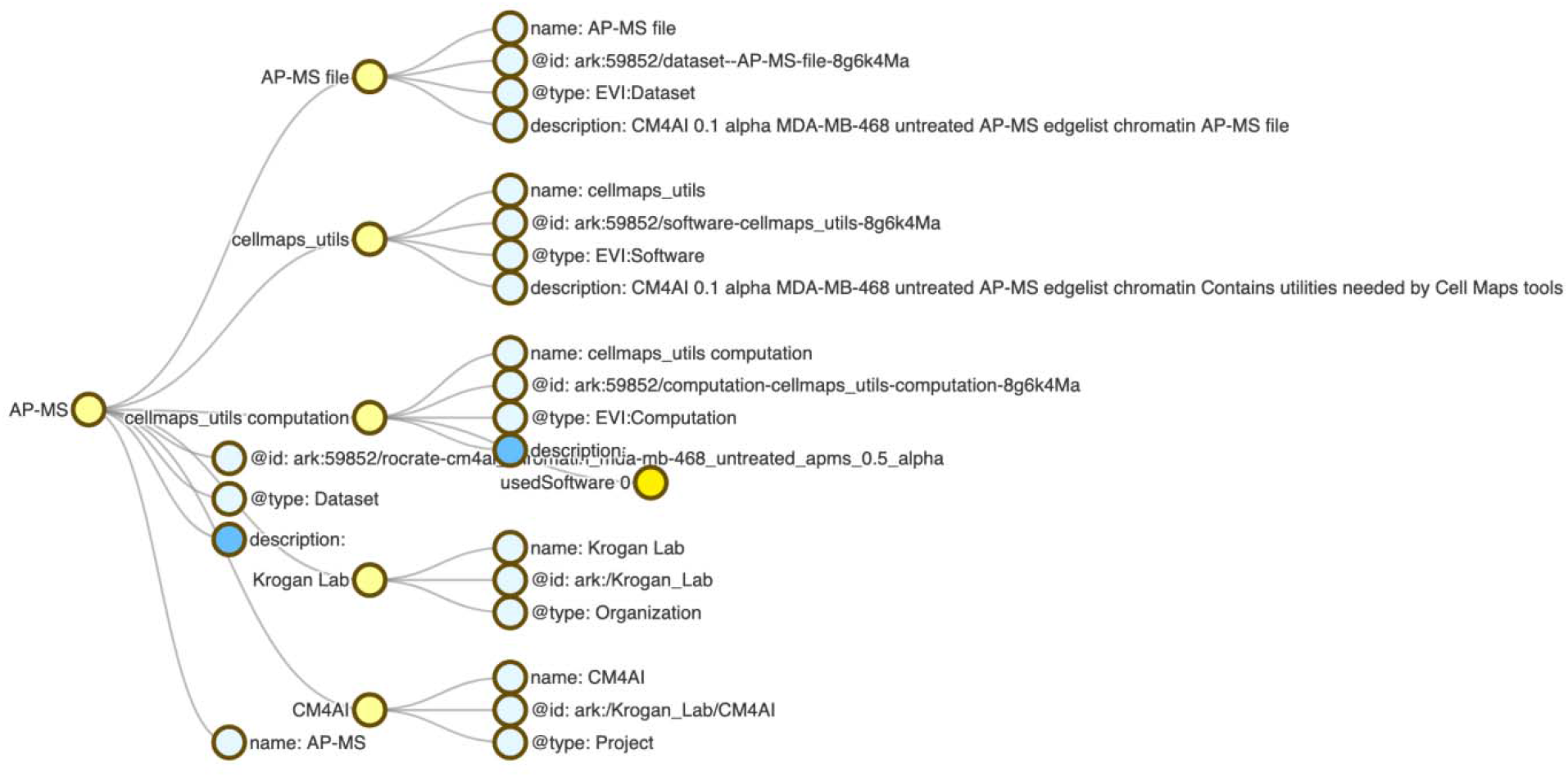
Evidence graph provenance visualization from human-readable landing page. This landing page is displayed on resolution of the RO-Crate’s ARK globally unique persistent ID. It was generated on an RO-Crate package from CM4AI’s 0.5 alpha data release. Landing page navigation tabs allow full metadata on the package to be displayed in human readable form, or alternatively in JSON-LD serialization.

Virginia’s LibraData data archive, an instance of Harvard’s Dataverse. Links to the archived RO-Crates are provided in the Data and Software Availability Statement.

PIDs generated by the server resolve to machine- and human-readable landing pages containing the metadata, expressed in the JSON-LD graph language using vocabularies from Schema.org, EVI, and other well-defined public ontologies. The complete RO-Crate dataset is attached as Supplemental Data.

Both the client-side and sever-side FAIRSCAPE packages are PIP-installable and are freely available, under the MIT open-source license (see Data and Software Availability Statement).

### 5. Teaming

The CM4AI project’s Teaming Module supports integration and expansion of technical / scientific expertise in CM4AI, emphasizing ethical considerations and diverse interdisciplinary perspectives. It facilitates effective communication and collaboration among CM4AI investigators and personnel, of varied geographic, disciplinary, and cultural backgrounds.

Teaming supports open dissemination of CM4AI-generated data, maps, and tools via the CM4AI web portal (http://www.cm4ai.org), using the U-BRITE platform^43^, to support open and trustworthy data management and sharing, ensuring the broad dissemination of the datasets, cell maps, and tools developed. This effort aligns with our commitment to FAIR principles and high-performance access, making our deliverables both accessible and useful to users.

Teaming has constructed an extensive scholarly network called a talent graph, based initially on publications of CM4AI investigators, which links to each person’s peer-reviewed research publications. Knowledge graphs involving this extended community of experts, publications, data sets, and tools are captured through this effort.

Collaboration with the NIH Common Fund Data Ecosystem (CFDE) for data curation helps integrate CM4AI contributions into the broader scientific community, and engagement with the Bridge2AI Center helps us identify novel channels for broadcasting our achievements, ensuring the outreach of our data, maps, and tools to those who can utilize them fully.

The Teaming module fosters a culture of open science and community involvement with mailing lists, shared online collaborating folders, shared documents, and remote messaging and collaboration software within CM4AI, across the Bridge2AI program, and in the broader scientific community, to promote an ethical, inclusive, and collaborative scientific environment. In collaboration with the Skills and Workforce Development module, it contributes significantly to enhancing the biomedical AI/ML workforce and fostering innovation in the field.

### 6. Skills and Workforce Development

The Skills and Workforce Development module works to enhance the biomedical AI/ML workforce by recruiting and training a diverse group of participants in the scientific approach, data standards, and tools that can be used to accelerate innovation from CM4AI. To recruit and prepare a biomedical workforce that is adept at both the data and life sciences, new training that will allow researchers to leverage the data sets produced by Bridge2AI and other programs is needed. To achieve this goal, we aim to broadly distribute the data and tools developed as part of the CM4AI project and provide broad training in these components. We are leveraging a multimodal approach to content delivery which includes asynchronous virtual training, hosted virtual events, and an in-person internship being co-hosted at two of our sites.

A key focus of the Skills and Workforce Development module is to cultivate a diverse biomedical AI/ML workforce. Our recruitment approach has included communication with investigators involved in the Bridge2AI program, integration with trainees at organizations involved with Bridge2AI, and community outreach to other academic organizations with an emphasis on those serving underrepresented communities. For our inaugural virtual CodeFest event, we provided training to a total of 38 registrants where 40% of the attendees identified as female and 30% came from underrepresented communities. We have also developed a Diversity, Equity, and Inclusion (DEI) Committee with representation from all Cores to direct initiatives which has at its core to partner with historically black colleges and universities (Meharry Medical and Morehouse School of Medicine) to increase the number of learners in the AI/ML field at all levels: graduate students: masters and PhD, postdoctoral, and junior investigators.

Our in-person internship will be hosted at Yale University and the University of California San Diego, with students jointly working on a final project between the two sites and supervised by faculty from both institutions. This internship will provide participants with the opportunity to learn 1) the impact of cell maps on biomedical research applications, 2) how to interact with the data and standards used by CM4AI, 3) how to develop a visible neural network using cell maps, and 4) how to integrate external data into an existing cell map for the ethical development of AI/ML applications.

### 7. Ethics

The Ethics team worked closely with Data Acquisition, Tools, and Standards Modules to develop a plan for ethical preparation, licensing, dissemination, and Data Access supervision of CM4AI datasets. This plan balances the openness of the data produced, with protection of the intellectual property, and supervision and monitoring of potential efforts at commercialization. In addition to CM4AI data governance initiatives, comprehensive guidelines and best practices are being established for the ethical development of AI systems leveraging this data. The Ethics team employs a methodology inspired by Value-Sensitive Design (VSD), facilitating the creation of normative standards and expectations aimed at ensuring the responsible design of datasets and AI technologies, in alignment with core values. This entails the Ethics team engaging in conceptual efforts to craft an axiological repository—a repository of values—to guide the articulation of design standards and expectations. Further advancing this work, the Ethics team is instrumental in formulating the CM4AI Life Cycle, a framework designed to elucidate governance milestones throughout the data generation and AI system development processes. This lifecycle framework underscores the commitment to embedding ethical considerations at every phase, ensuring accountability and value alignment from inception through deployment.

Recognizing the importance of diverse perspectives in shaping ethical guidelines, the Ethics team is dedicated to conducting mixed empirical research, combining qualitative and quantitative methods. This research is critical for capturing a broad spectrum of community insights, thus enriching the development of guidelines and best practices with a multifaceted understanding of stakeholder values and concerns. Moreover, the team is set to develop a suite of tools aimed at enhancing awareness and comprehension of the ethical, legal, and social ramifications associated with CM4AI data and the consequent AI systems. These tools are envisioned to bolster ethical decision-making, providing vital support for navigating the complex landscape of AI ethics. This comprehensive approach not only aims to elevate the standards of AI development within CM4AI but also seeks to contribute to the broader discourse on responsible AI, promoting a culture of ethical integrity, inclusivity, and societal benefit.

## Results and Progress to Date

The CM4AI project has made significant progress with efforts by each of our six modules (Data, Tools, Standards, Ethics, Skills, Teaming) relevant to each of the three program pillars: Data – composed of the Data, Tools, and Standards Modules; People – composed of the Skills and Teaming Modules; and Ethics – the Ethics Module.

The Data Module completed parallel maps of protein-protein interactions, protein subcellular distributions, and transcriptional states associated with a near-comprehensive set of chromatin regulators encoded by the human genome (Pillar: Data). All of these data were generated in ethically sourced breast cancer cells (Pillar: Ethics) and will be extended to ethically sourced human induced pluripotent stem cells (iPS) in Year 2.

The Tools Module developed an early (alpha) release of the Multi-Scale Integrated Cell (MuSIC) toolkit for assembling large-scale maps of human cells from fusion of protein interaction and imaging data, including a highly automated pipeline that integrates with Standards FAIRSCAPE toolkits. We expect to submit a mature version for publication as a Protocols paper in early fall

The Standards Module established FAIR AI-Readiness metadata and validation toolkits, which have been used to package and release our cell-line perturbation and measurement data.

The Ethics Pillar established guidelines addressing socio-ethical challenges, positioning us to now embark on stakeholder engagement and explainable AI development. Preliminary work has been done to develop a first version of the Value Repository to support the establishment of these guidelines^44,45^. The Ethics Pillar also supported the work initiated by the Bridge2AI Ethics Working Group for supporting the Bridge2AI Data Sharing and Dissemination Working Group in developing a Code of Conduct. The Code of Conduct outlines basic requirements for participants in the Bridge2AI Open House to access B2AI data (including CM4AI datasets), to attest and commit to it prior to accessing the data.

The Workforce Development Module identified gaps in training of the functional genomic and biomedical AI/ML workforce; and led the organization of a highly successful CodeFest in March 2024, in collaboration with the Tools, Standards, Data, and Teaming Modules. The CodeFest was attended by both within-project and external participants. Our Workforce Development Module is now transitioning towards collaborative content creation and delivery (Pillar: People).

Finally, the Teaming Module facilitated cross-disciplinary collaboration, completed the open-access CM4AI web portal, and initiated a Diversity Equity and Inclusion (DEI) Committee (Pillar: People). Beyond these specific activities, CM4AI has been actively engaged in activities of the greater Bridge2AI Consortium, including a face-to-face meeting in Pentagon City, Virginia; the overall Bridge2AI Steering Committee and workgroups; and submission of numerous auxiliary concepts for supplemental funding and a U-24 submission for cross-cutting integration across projects in the NIH Common Fund Data Ecosystem.

Initial Datasets, Tools and Standards associated with all three parallel mapping platforms have been released and are publicly available at our CM4AI web portal^30^. These data were also made available at our CodeFest in March 2024, accompanied by detailed tutorials explaining how the data were derived and integrated, and providing production ready software for producing the datasets. This first data release was made possible by intensive cross-module collaboration by dozens of staff as well as end-to-end operation of all aspects of our CM4AI platform, from data generation to data analysis by advanced toolsets to FAIR-compliant ethical AI-readiness standards for data access and distribution.

Data packages from CM4AI including links to software used in preparing the data, input datasets, dataset schemas, and deep provenance graphs, are available on the CM4AI web portal and will soon be available on the Dataverse NIH approved generalist repository.

All CM4AI integration and packaging software is freely available and licensed under nonrestrictive open-source licenses. Data packages are licensed under Creative Commons CC-BY-NC-SA 4.0 license terms^31^, which include a requirement for attribution to the dataset authors and the copyright holding institutions, and citation of this article. Commercial use requires a separate license negotiation with the copyright holder (UCSD, Stanford, and/or UCSF depending upon the specific data package in question). A Data Access Committee will supervise ethical matters related to dataset distribution and potential dual licensing for commercial use.

## Conclusions

The Bridge2AI Functional Genomics project, “Cell Maps for Artificial Intelligence” (CM4AI), is a continuing effort which has already published valuable and re-usable datasets and software, advanced the notions of ethical AI-readiness, developed a strong teaming approach within the project including a shared portal, and provided hands-on training to the research community in use of these tools.

This article presents the work as a whole in preprint form and will be continuously updated as CM4AI progresses, with citations to detailed publications in each contributing area.

With these measures, it is our intention that CM4AI become a highly transformative platform for ethical, explainable, and interpretable biomedical AI, deepen our understanding of processes of human disease and health, help to train a new generation of researchers, enable development of novel cures, and assist researchers and clinicians in equitably improving human lives.

## Data and Software Availability Statement

The most recent data and metadata produced by CM4AI are licensed for reuse under the Creative Commons Attribution Non-Commercial Share-Alike International 4.0 License (https://creativecommons.org/licenses/by-nc-sa/4.0/) and are available in LibraData, the University of Virginia’s archival data repository:

*Clark T, Mohan J, Schaffer L, Obernier K, et al. Cell Maps for* Artificial Intelligence - Data Release”, https://doi.org/10.18130/V3/DXWOS5, University of Virginia Dataverse, V1

LibraData is the University of Virginia’s instance of Dataverse, an NIH-approved generalist repository.

*Attribution requirements for these datasets include attribution to the copyright holders and the Cell Maps for Artificial Intelligence project, as referenced in the datasets, and citation of the present article:*

*Clark T, Mohan J, Schaffer L, Obernier K, Al Manir S, Churas CP, Dailamy A, Doctor Y, Forget A, Hansen JN, Hu M, Lenkiewicz J, Levinson MA, Marquez C, Nourreddine S, Niestroy J, Pratt D, Qian G, Thaker S, Bélisle-Pipon J-C, Brandt C, Chen J, Ding Y, Fodeh S, Krogan N, Lundberg E, Mali P, Payne-Foster P, Ratcliffe S, Ravitsky V, Sali A, Schulz W, Ideker T. Cell Maps for Artificial Intelligence: AI-Ready Maps of Human Cell Architecture from Disease-Relevant Cell Lines. BioRXiv, May 2024*.

Copyright © 2024 to these datasets is held by the Regents of the University of California, except where otherwise indicated. Raw image data for spatial proteomics is copyright © 2024 by The Board of Trustees of the Leland Stanford Junior University.

Software comprising the Tools data integration pipeline is available under BSD-3 open source license in GitHub (for alpha level tools) or in the Zenodo long-term archive (for production-ready tools). Links to these tools are packaged in RO-Crates as part of CM4AI provenance representations.

The FAIRSCAPE AI-readiness framework is described with a tutorial and instructions for installation at https://fairscape.github.io. Copyright © 2024 to this software is held by the Rector and Board of Visitors of the University of Virginia. It is available under open source MIT License.

The open source *Integrative Modeling Platform* (IMP) package is available at http://integrativemodeling.org.

Software packages will be versioned as the project progresses, and the version used to produce each dataset will be referenced in the dataset’s metadata.

## Acknowledgements

This work was funded by the National Institutes of Health under awards 1OT2OD032742-01 (Bridge2AI Functional Genomics) and 5U54HG012513-02 (Bridge2AI Bridge Center), and by the Frederick Thomas Fund of the University of Virginia.

We also acknowledge the helpful assistance of members of the other Bridge2AI components, which helped to make this work possible, including but not limited to the Bridge2AI Standards Working Group; the Bridge2AI Data Sharing and Dissemination Working Group; and members of the Bridge2AI CHoRUS Critical Care Medicine Data Generation Project.

