## Supplemental Data File for "Cell Maps for Artificial Intelligence: AI-Ready Maps of Human Cell Architecture from Disease-Relevant Cell Lines": readme.txt

Affinity Purification Mass spectrometry (AP-MS)
-----------------------------------------------

Affinity Purification-Mass Spectrometry (AP-MS) is a powerful technique used to uncover the intricate networks of
protein interactions within cells. Different methods can be used but the most popular approach is to attach a purifiable
tag to a protein of interest, the "bait". In the context of CM4AI the tag is directly attached to the endogenous protein
of interest using gene editing to maintain physiological level of expression. The tagged bait is then purified from cell
extract with its interacting partners. Once separated, the captured proteins are identified using a mass spectrometer
allowing to infer protein-protein interactions. Understanding these interactions allows can unravel complex cellular
processes, discover new roles for proteins, and gain insights into diseases caused by disrupted protein networks.


Data processing
---------------

Raw data acquired on a Orbitrap Fusion™ Lumos™ Tribrid™ Mass Spectrometer (Thermo Scientific)
Raw data are processed using Maxquant v1.6.12.0 using default parameter with LFQ quantification and the
“202209_uniprot_human_reviewed.fasta” file as reference.
Evidence file from maxquant are processed using ArtMS in R to generate SAINT input files using the
following command lines. SAINTexpress was run locally using version SAINTexpress_v3.6.3
SAINT is designed to assess the reliability and significance of protein-protein interactions identified
through mass spectrometry experiments. It takes into account both the presence and abundance of proteins
in different experimental conditions or replicates. The method uses a probabilistic model to estimate the
probability that a given interaction is a true positive rather than a random or non-specific interaction.
This approach helps to distinguish true interactions from noise in large-scale protein interaction datasets.

apms.tsv column labels
----------------------

Bait: Name of the pull downed protein

Prey: Uniprot ID number of identified proteins by MS in pull down (putative bait interactor).

PreyGene.x: Uniprot protein name of identified protein by MS in pull down (putative bait interactor).

Spec: Number of spectral count in each test biological replicates (separated by | ).

SpecSum: Sum of Spectral counts in test samples.

AvgSpec: Average Spectral counts across replicates in test samples.

NumReplicates.x: Number of replicates in test samples.

ctrlCounts: Number of spectral count in each control replicates (separated by | ).

AvgP.x: Average probability that an interaction is true, measure of the likelihood that a given
        interaction is a true positive rather than a random or non-specific interaction. A lower AvgP indicates
        higher confidence in the interaction being genuine.

MaxP.x: maximum probability associated with a protein interaction in the context of its prey-bait pair.
        Similar to AvgP, a lower MaxP suggests a higher likelihood of the interaction being true.

TopoAvgP.x: extension of the AvgP score that also takes into consideration the topology of the
            interaction network. It incorporates information about the hierarchical structure of the interaction
            data to provide a refined assessment of the interactions.

TopoMaxP.x: topology-aware score that considers the maximum probability of an interaction in the
            context of the interaction network's topology.

SaintScore.x: composite score that integrates multiple aspects of the interaction data, including
              spectral counts and probability estimates. It's designed to prioritize interactions based on their
              strength and reliability. Higher SaintScores indicate interactions that are more likely to be true.

logOddsScore: Logarithm of the odds ratio between test and control conditions for each prey as a
              measure of interaction significance. The LogOddsScore is a statistical score that represents the
              logarithm of the odds ratio for a protein-protein interaction. It's used to quantify the strength
              and significance of the association between two proteins in an interaction network. The odds ratio
              compares the likelihood of the interaction occurring to the likelihood of it not occurring. Taking
              the logarithm of the odds ratio often helps to transform the score into a more symmetric and interpretable
              form, making it easier to compare and analyze the interactions. Higher LogOddsScores typically indicate
              stronger evidence for the interaction.

FoldChange.x: represents the ratio of the abundance of a protein or interaction in one experimental
              condition (Test) compared to another (control). It helps assess whether the abundance of a protein
              changes significantly between different conditions.

BFDR.x: Bayesian False Discovery Rate


SAINT References:
https://saint-apms.sourceforge.net/Main.html
https://www.ncbi.nlm.nih.gov/pmc/articles/PMC4102138/
https://pubmed.ncbi.nlm.nih.gov/21131968/
